# Improved whole-mount immunofluorescence protocol for consistent and robust labeling of adult *Drosophila melanogaster* adipose tissue

**DOI:** 10.1101/2024.04.12.589269

**Authors:** Rachael K. Ott, Alissa R. Armstrong

**Affiliations:** Department of Biological Sciences University of South Carolina, Columbia

**Keywords:** *Drosophila*, adult fat body, immunofluorescence

## Abstract

Energy storage and endocrine functions of the *Drosophila* fat body make it an excellent model for elucidating mechanisms that underlie physiological and pathophysiological organismal metabolism. Combined with *Drosophila’s* robust genetic and immunofluorescence microscopy toolkits, studies of *Drosophila* fat body function are ripe for cell biological analysis. Unlike the larval fat body, which is easily removed as a single, cohesive sheet of tissue, isolating intact adult fat body proves to be more challenging, thus hindering consistent immunofluorescence labeling even within a single piece of adipose tissue. Here, we describe an improved approach to handling *Drosophila* abdomens that ensures full access of the adult fat body to solutions generally used in immunofluorescence labeling protocols. In addition, we assess the quality of fluorescence reporter expression and antibody immunoreactivity in response to variations in fixative type, fixation incubation time, and detergent used for cellular permeabilization. Overall, we provide several recommendations for steps in a whole mount staining protocol that results in consistent and robust immunofluorescence labeling of the adult *Drosophila* fat body.

**SUMMARY STATEMENT:** Optimization of adult *Drosophila* fat body fluorescence microscopy via modifications of tissue handling, fixation, and permeabilization steps in a whole mount immunolabeling protocol.

## INTRODUCTION

From insects to mammals, adipose tissue serves as an energy storage and endocrine organ (Adamczak and Wiecek, 2013; Meschi and Delanoue, 2021; Navarro-Perez et al., 2023), controlling several aspects of organismal physiology, such as lipid metabolism, reproduction, blood pressure regulation, thermoregulation, immune system modulation, and insulin sensitivity (Adamczak and Wiecek, 2013). Adipose tissue dysfunction results from obesity and is associated with pathophysiologies including dyslipidemia, infertility, high blood pressure, chronic inflammation, several cancers, and type 2 diabetes (Longo et al., 2019). For example, diet-induced obesity leads to adipocyte hypertrophy and hyperplasia, as well as altered adipokine profiles and increased secretion of pro-inflammatory cytokines (Park et al., 2014). Given that adipose tissue acts as an essential mediator of energy homeostasis in both health and disease, it is critical to gain a better understanding of the cellular and molecular mechanisms that allow adipocytes to relay physiological information to peripheral organs. The incredible genetic toolkit (Caygill and Brand, 2016; Fölsz et al., 2022; Gratz et al., 2023), presence of organ systems analogous to mammals, and high degree of genetic conservation (Ugur et al., 2016) make *Drosophila melanogaster* an excellent *in vivo* model organism to address the role that adipose tissue plays in inter-organ communication.

Derived from mesoderm, the larval fat body exists throughout the anterior-posterior axis as a sheet of tightly adhered polygonal adipocytes that dissociate into individual spherical cells at the larval-to-pupal transition (Zheng et al., 2016). During the pupal-to-adult transistion, adipocyte precursors migrate from the thorax to the abdomen, proliferate, and undergo fusion to generate the adult fat body (Tsuyama et al., 2023). While the vast majority of fat body mass is located in the abdomen, adipose tissue is also found in the head (pericerebral fat) and thorax (Roman et al., 2001). In adult flies, adipocytes, the major cellular component, and hepatocyte-like oenocytes make up the abdominal fat body. Like mammalian adipose tissue, the *Drosophila* fat body has energy storage and endocrine roles (Parra-Peralbo et al., 2021). The activity of triacylglycerol synthesis enzymes, lipid storage proteins, and lipases in the fat body regulates the balance between energy storage and mobilization, particularly in response to nutrient availability (Heier et al., 2021; Li et al., 2019). During larval development and adulthood, the *Drosophila* fat body plays a major role in controlling cell, tissue, and organismal size (Lim et al., 2019; Manière et al., 2020; Schmitt et al., 2015; Ugrankar-Banerjee et al., 2023), the humoral immune response (Bland, 2023), and inter-organ communication (Armstrong and Drummond-Barbosa, 2018; Meschi and Delanoue, 2021; Weaver and Drummond-Barbosa, 2019).

*Drosophila* serves as an excellent model for adipose tissue biology as well as obesity and metabolic disorders (Musselman and Kühnlein, 2018). Several well-established genetic, biochemical, molecular, and histological methods have uncovered key features of fat body development, function, and dysfunction (Nayak and Mishra, 2019). Whole-mount fluorescence immunohistochemistry has been instrumental in assessing morphology and protein localization within cells of a given tissue. Here, we describe an improved protocol for whole-mount immunofluorescence of the abdominal fat body in *Drosophila* adults.

## RESULTS

### Abdominal carcass stabilization enhances adult fat body immunostaining

In our previous approach to whole-mount adult fat body immunostaining, we incubated isolated abdominal carcasses, excluding reproductive and gut tissues, in solutions of the immunofluorescence protocol suspended in microfuge tubes. This method often results in inconsistent immunolabeling quality. For example, we observed robust, weak, and undetected labeling of fat body tissue from abdominal carcasses carried through the process in the same tubes. We reasoned that this labeling variability resulted from solution inaccessibility due to 1) ventral abdominal flaps covering the internal dorsal abdomen, and 2) abdominal carcasses becoming lodged into microfuge tube caps. To provide continual fat body tissue access to solutions, we have developed a method in which abdominal carcasses are carried through the immunostaining process while being pinned open and stabilized (**Fig. 1**). For dissection, an adult fly is submerged in an appropriate buffer (Grace’s medium, insect medium, or 1X PBS), and forceps are used to remove the abdomen from the thorax and head (**Fig. 1A**). Forceps were used to gently tear along the anterior-posterior axis of the ventral abdomen and remove the reproductive tissues and gut (**Fig. 1B**). After removing the last one or two abdominal segments (**Fig. 1C**) abdominal carcasses were transferred to a well in a Sylgard^TM^-coated, 6-well tissue culture dish filled with dissecting media. Since we housed multiple abdominal carcasses in each well, each was strategically placed to maximize the spacing. While gently pressing an abdominal carcass to the coated surface as a counterpressure, forceps were used to pin down the first “corner” with a 0.10 mm Austerlitz insect pin (**Fig. 1D**) followed by pinning the remaining edges of the abdominal carcass to expose the internal cavity (**Fig. 1E**). This process was repeated for up to six abdominal carcasses as needed (**Fig. 1F**) (the vertical extension of pins out of the wells makes it more difficult to pin more). Fixation, washes, and antibody incubations were performed in these wells by carefully removing and replacing each solution. After completing the immunostaining process (described in the Materials and Methods), insect pins were gently removed with forceps, and abdominal carcasses were transferred to microfuge tubes containing mounting media. To mount fat body tissue, a glass Pasteur pipet was used to transfer abdominal carcasses to a droplet of mounting media on a microscope slide followed by carefully separating fat body tissue from the cuticle using forceps and a needle (**Fig. 1G**). Cuticles were discarded prior to applying the coverslip.

**Fig. 1.**
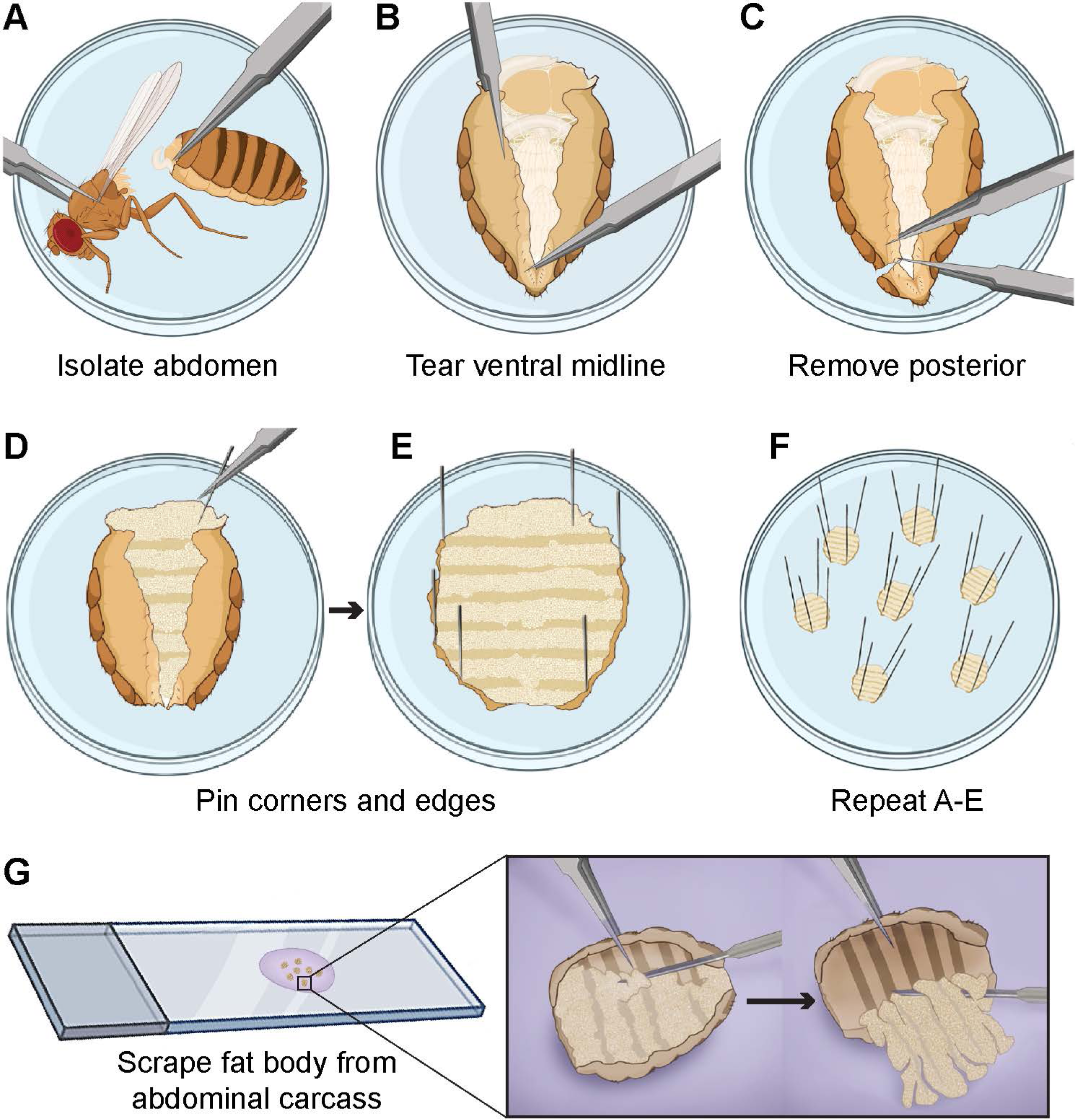
Pinned open abdominal carcasses for whole-mount adult fat body immunofluorescence. Forceps used to (A) remove the thorax and head from abdomens, (B,C) tear along the ventral midline to open the abdominal cavity, and remove reproductive and gut tissues. (D-F) Insect pins to secure abdominal carcass edges onto Sylgard®-coated wells of a 6-well tissue culture plate. Upon completion of the immunostaining protocol, (G) insect pins carefully removed and abdominal carcasses transferred to a slide in droplet of mounting media. Forceps and needles used to separate the fat body from the exoskeleton, which is removed prior to coverslip attachment. Created with BioRender.com.

### Paraformaldehyde fixation provides optimal preservation of adipocyte cellular morphology

Aldehyde- and solvent-based fixatives are widely used to preserve tissue and cellular structure for use in immunohistochemistry and immunocytochemistry approaches. Aldehyde-based fixatives, such as paraformaldehyde (PFA) and glutaraldehyde (GA), crosslink proteins together while solvent-based fixatives, such as methanol and acetone, dehydrate and precipitate proteins to preserve cellular morphology (Hobro and Smith, 2017). Paraformaldehyde fixation has long been the standard for immunofluorescence studies while glutaraldehyde fixation is traditionally used in electron microscopy (Foissner and Hoeftberger, 2019; Im et al., 2019). Methanol and acetone are popular among *in vitro* immunostaining protocols (Jamur and Oliver, 2010). To determine the best fixation approach that provides optimal preservation of *Drosophila* adult fat body for use in immunofluorescence studies, we evaluated fixative type, concentration, and incubation time using the abdominal carcass pinning technique described above.

To avoid confounding effects of variable antibody immunoreactivity, the initial test of fixatives made use of fat body samples from two transgenic fly lines known to control expression in the adult fat body: *cg-Gal4; UAS-myr-RFP*, that drives expression of red fluorescent protein (RFP) at the cellular membrane, and *FB-Gal4; UAS-GFP.nls,* that drives expression of green fluorescent protein (GFP) in the nucleus (Armstrong et al., 2014; Grönke et al., 2003; Pastor-Pareja and Xu, 2011). We first tested methanol and acetone fixation and observed very poor fat body tissue preservation (data not shown). Next, we asked if paraformaldehyde and glutaraldehyde fixation provide equivalent preservation of adult fat body tissue. Adipocytes from *FB-Gal4; UAS-GFP.nls* flies fixed in paraformaldehyde showed diffuse cytoplasmic GFP with enriched nuclear GFP expression (**Fig. 2A**). Similarly, adipocytes from *cg-Gal4, UAS-myr-RFP* flies showed robust membrane RFP labeling (**Fig. 2B**). In contrast, fluorescence in these cellular structures was not preserved in adipocytes from flies fixed in glutaraldehyde. With 4% glutaraldehyde fixation, *FB-Gal4; UAS-GFP.nls* samples did not show GFP nuclear enrichment (**Fig. 2C**) and *cg-Gal4, UAS-myr-RFP* samples had perinuclear RFP aggregates instead of membrane-restricted labeling (**Fig. 2D**). At a much lower glutaraldehyde concentration, nuclear GFP expression was restored (**Fig. 2E**) while membrane labeling was minimal (**Fig. 2F**); however, tissue preservation was still poor. We also assessed whether tandem aldehyde fixation might enhance adult fat body preservation by comparing varying percentages of initial paraformaldehyde fixation followed by post-fixation with glutaraldehyde in the following combinations: 2.5% PFA, 2.5% GA, 2.5% PFA, 0.1% GA, and 2.5% PFA, 0.05% GA. Of these combinations, 2.5% PFA followed by 0.05% GA showed improved tissue preservation and fluorescence reporter detection compared to glutaraldehyde fixation alone (**Fig. 2G, H compared to Fig. 2C-F**). Using this approach, nuclear GFP labeling in *FB-Gal4; UAS-GFP.nls* adipose tissue was comparable to paraformaldehyde fixation (compare **Fig. 2G** and **2A**). By contrast, tandem fixation provided inconsistent preservation of myristoylated RFP at the cell membrane in *cg-Gal4* samples and retained prominent cytoplasmic signal from perinuclear puncta (**Fig. 2H**). Taken together, these results suggest that not all cross-linking aldehyde fixatives provide equivalent preservation of adipocyte morphology for immunofluorescence studies in the adult fat body and that paraformaldehyde is the fixative of choice.

**Fig. 2.**
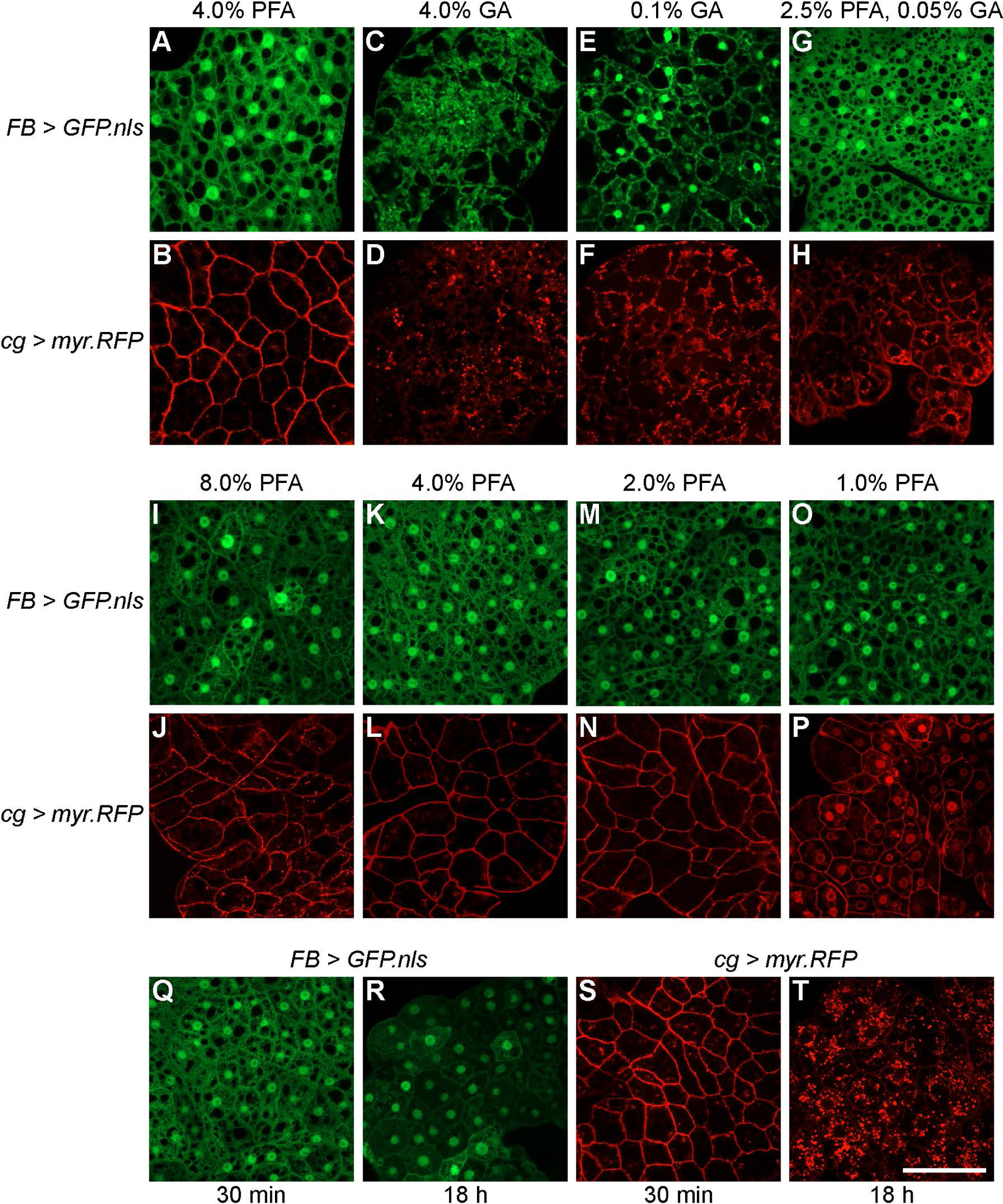
Comparison of fixative type, concentration, and time for adult fat body preservation. (A,B) Fat body tissue fixed with 4% paraformaldehyde alone, (C,D) 4% glutaraldehyde alone, (E,F) 0.1% glutaraldehyde alone, and (G,H) 2.5% paraformaldehyde/0.05% glutaraldehyde in combination. Fat body samples from (I-O) *FB-Gal4 > UAS-GFP.nls* (green) and (J-P) *cg-Gal4 > UAS-myr.RFP* (red) transgenic lines fixed with decreasing concentrations of paraformaldehyde (8%, 4%, 2%, and 1%) for 30 minutes. Fat body samples from (Q,R) *FB-Gal4 > UAS-GFP.nls* and (S,T) *cg-Gal4 > UAS-myr.RFP* transgenic flies fixed with 4% paraformaldehyde for 30 minutes or 18 hours. Scale bar = 50 μm

Effective paraformaldehyde fixation relies on the extent of adduct formation between cellular components (Hobro and Smith, 2017), prompting us to ask if there is an optimal paraformaldehyde concentration that preserves adult adipose tissue with a 30-minute fixation time. At all paraformaldehyde concentrations tested, 8%, 4%, 2%, and 1%, *FB-Gal4; UAS-GFP.nls* samples showed enriched nuclear GFP labeling (**Fig. 2I, K, M, O**). Similarly, *cg-Gal4, UAS-myr-RFP* samples across all tested paraformaldehyde concentrations showed RFP at the cell membrane (**Fig. 2J, L, N, P**). However, at the lowest concentration, 1% paraformaldehyde, we observed prominent nuclear RFP localization in addition to the membrane fluorescence (**Fig. 2P**). These data suggest that, while a lower limit exists, a range of paraformaldehyde concentrations provides robust preservation of the adult fat body within a 30-minute fixation period.

The availability of molecular substrates in the cytosol influences paraformaldehyde fixation efficiency. Despite its rapid penetration, paraformaldehyde reaction rates with different proteins depend on protein amino acid composition (Kamps et al., 2019) and tertiary structure (Hoffman et al., 2015). On one hand, protein stabilization via initial adduct formation is easily reversed while excessive cross-linking can hinder immunoreactivity by epitope masking (Kiernan, 2015). Therefore, preservation of sample integrity and immunoreactivity require an appropriate fixation time. We evaluated adipocyte morphology and reporter detection following 4% paraformaldehyde fixation for 30 minutes, 1 hour, 3 hours, and 18 hours. Fixation up to 18 hours did not significantly alter adipocyte morphology or GFP expression patterns in *FB-Gal4; UAS-GFP.nls* samples, though nuclear fluorescence intensity was slightly reduced (compare **Fig. 2R** and **2Q**). By contrast, 18-hour fixation of adipocytes from *cg-Gal4, UAS-myr-RFP* samples contained dense cytoplasmic and perinuclear enriched RFP puncta and undetectable membrane labeling (**Fig. 2T**), unlike the robust membrane RFP in the samples fixed for 30 minutes (**Fig. 2S**). Similarly, fixation for one or three hours was not compatible with either transgenic reporter as evidenced by altered expression patterns, i.e., prominent membrane labeling in *FB-Gal4, UAS-GFP.nls* samples and cytoplasmic puncta without membrane labeling in *cg-Gal4; UAS-myr-RFP* samples (data not shown). Collectively, these results demonstrate that prolonged paraformaldehyde fixation negatively impacts fluorescence reporter activity in adult adipose tissue and that 30 minutes is an ideal fixation time.

### Triton X-100 and Tween 20 allow for robust immunolabeling of adipocyte cellular components

In tissues fixed by cross-linking agents, such as paraformaldehyde, antibody access to intracellular epitopes often requires membrane permeabilization. Immunofluorescence protocols often employ detergents, such as Triton X-100 and Tween 20, as permeabilization reagents. Their amphiphilic character and ability to intercalate between membrane phospholipids (Banfalvi, 2016; Mattei et al., 2017) make these nonionic detergents ideal for cell membrane permeabilization. However, detergent-based permeabilization can be deleterious to cellular structures at high concentrations (Goldenthal et al., 1985) and can affect the overall antibody distribution within the cell (Whelan and Bell, 2015). Since robust immunolabeling relies on optimal detergent use, we assessed immunostaining quality in the adult fat body treated with low (0.1%) and high (0.5%) concentrations of Triton X-100 and Tween 20. We labeled adipocyte cellular membranes with alpha spectrin and DE-cadherin antibodies and cytoskeletal elements with a tubulin antibody and F-actin dye (phalloidin). For all conditions tested, we observe comparable membrane labeling with the alpha spectrin antibody (**Fig. 3A-D**). Fat body samples labeled with DE-cadherin antibody and permeabilized with 0.1% Tween 20 showed labeling restricted to the membrane (**Fig. 3F**) while 0.5% Tween 20 showed labeling primarily at the membrane but also in cytoplasmic puncta (**Fig. 3H**). While permeabilization with Triton X-100 resulted in enriched labeling of DE-cadherin at the membrane, we noticed that the labeling was more punctate and there was diffuse cytoplasmic labeling not observed when samples were permeabilized with Tween-20 (**Fig. E, G**). Similar to labeling with the alpha spectrin antibody, phalloidin and tubulin labeling had comparable immunoreactivity across all conditions (**Fig. 3I-P**). Altogether, this data indicates that 0.1% Tween 20 provides specific and robust labeling across all antibodies tested. In addition, the slight variation in labeling quality suggests that antibody immunoreactivity should be assessed with different detergents and concentrations to identify the optimal detergent and concentration.

**Fig. 3.**
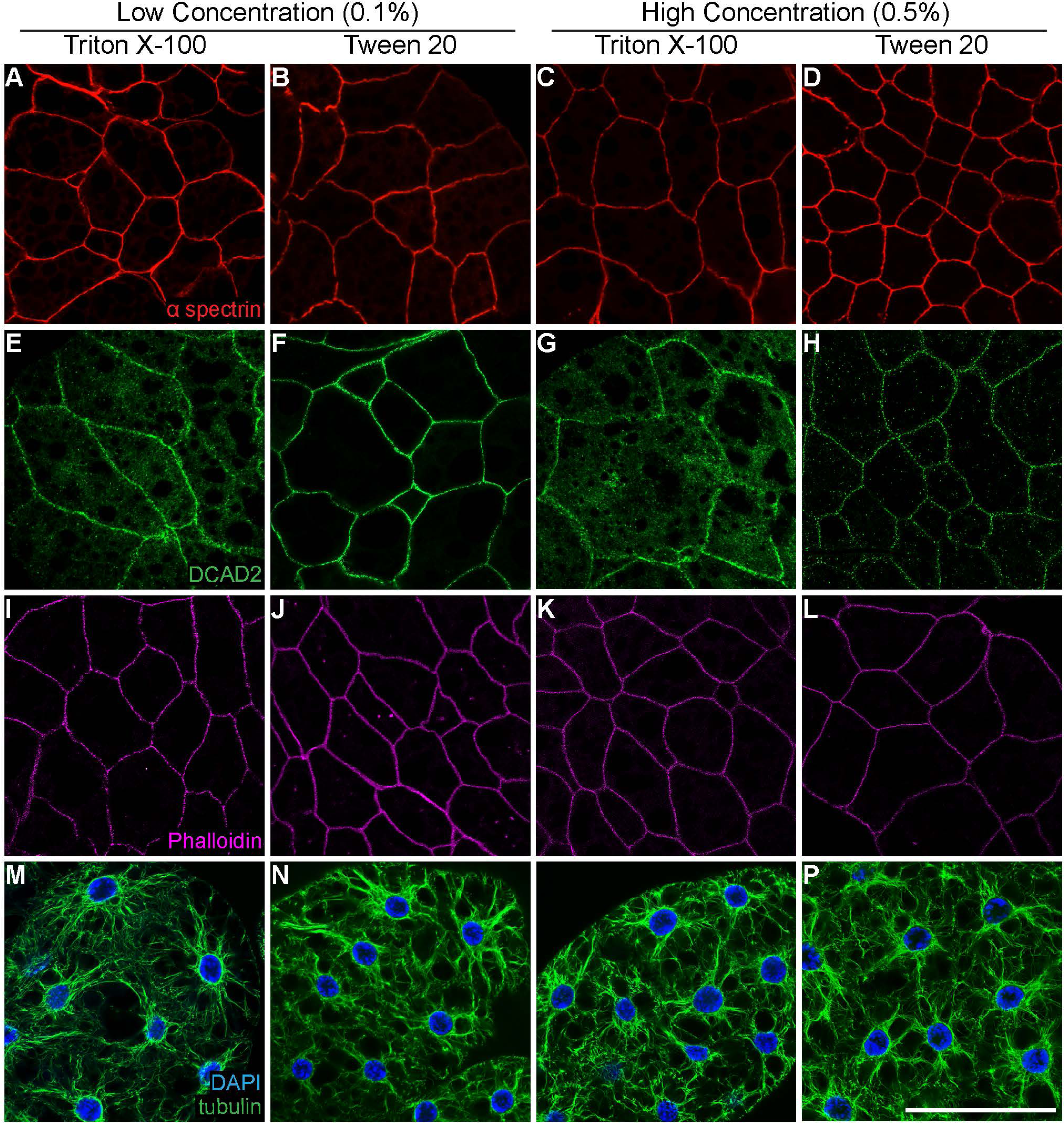
Comparison of detergent type and concentration for adult fat body permeabilization. Adult fat body tissue from *D. melanogaster* females immunolabeled with (A-D) anti-alpha spectrin (red), (E-H) anti-DE-Cadherin (green), (I-L) phalloidin (magenta), and (M-P) anti-alpha tubulin (green) (DAPI, blue) using either high (0.5%) or low (0.1%) concentrations of Triton X-100 or Tween-20 as the detergent in wash and blocking solutions. Scale bar = 50 μm

### Visualization of lipid droplets with lipophilic dyes Nile Red and BODIPY

From flies to humans, intracellular lipid droplets support many of the specialized functions of adipose tissue, including energy storage (Olzmann and Carvalho, 2019). *Drosophila* adipocytes contain many lipid droplets that vary in size (small to large) and location within the cell (peripheral and medial). Fluctuations in lipid droplet size reflect organismal physiology and demands for energy expenditure. For example, adipocytes in female flies fed a protein-poor diet show reductions in the number and size of lipid droplets (Matsuoka et al., 2017). The lipophilic fluorophores Nile Red and BODIPY are routinely used to detect lipid droplets in adipocytes (Armstrong et al., 2014) and other cell types in tissues with high metabolic demands, such as the *Drosophila* ovary (Giedt et al., 2023), gut (Nayak and Mishra, 2021), and brain (Liu et al., 2015). The high extinction coefficients and fluorescence quantum yields of Nile Red and BODIPY (Greenspan and Fowler, 1985; Loudet and Burgess, 2007) make these dyes convenient for use in multi-labeling fluorescence microscopy. However, for Nile Red, the wide range of excitation and emission spectra and the dependence of fluorescence intensity on solvent (Greenspan et al., 1985) makes it important to determine the appropriate conditions for specific lipid droplet labeling.

We evaluated several concentrations of Nile Red and BODIPY 505/515 dyes using the dissection and immunostaining procedure described above. At 2 μg/ml, 1 μg/ml, and 35 ng/ml concentrations of Nile Red, lipid droplet labeling was detected in both the red and green channels, which could not be mitigated by changing solvent (1X PBS, dH20, or 50% glycerol in 1X PBS) or exposure time (1 hr, 45, 40, 30, or 20 minutes) (data not shown). However, abdominal carcasses incubated in 25 ng/ml Nile Red diluted in 50% glycerol for 30 minutes showed lipid droplet fluorescence only in the red-light visible range (**Fig. 4A**). Similarly, samples incubated in 25 ng/ml BODIPY 505/515 in PBS for 30 minutes showed lipid droplet fluorescence only in the green-light visible range (**Fig. 4B**). At concentrations above 25 ng/ml (50 ng/ml, 250 ng/ml, 500 ng/ml, 1 μg/ml, 2 μg/ml), the extremely high BODIPY fluorescence intensity levels made it impossible to acquire images without over saturation (data not shown). Nile Red and BODIPY were compatible with co-labeling as evidence by anti-alpha spectrin immunostaining at adipocyte plasma membranes (**Fig. 4A, B**). Nile Red and BODIPY labeled the wide range of lipid droplet sizes from very small to very large. BODIPY was particularly efficient at detecting small cortical lipid droplets at the cell periphery (**Fig. 4C**). Thus, relatively low concentrations of both lipid dyes and short incubation times allow for robust detection of lipid droplets in adult adipocytes.

**Fig. 4.**
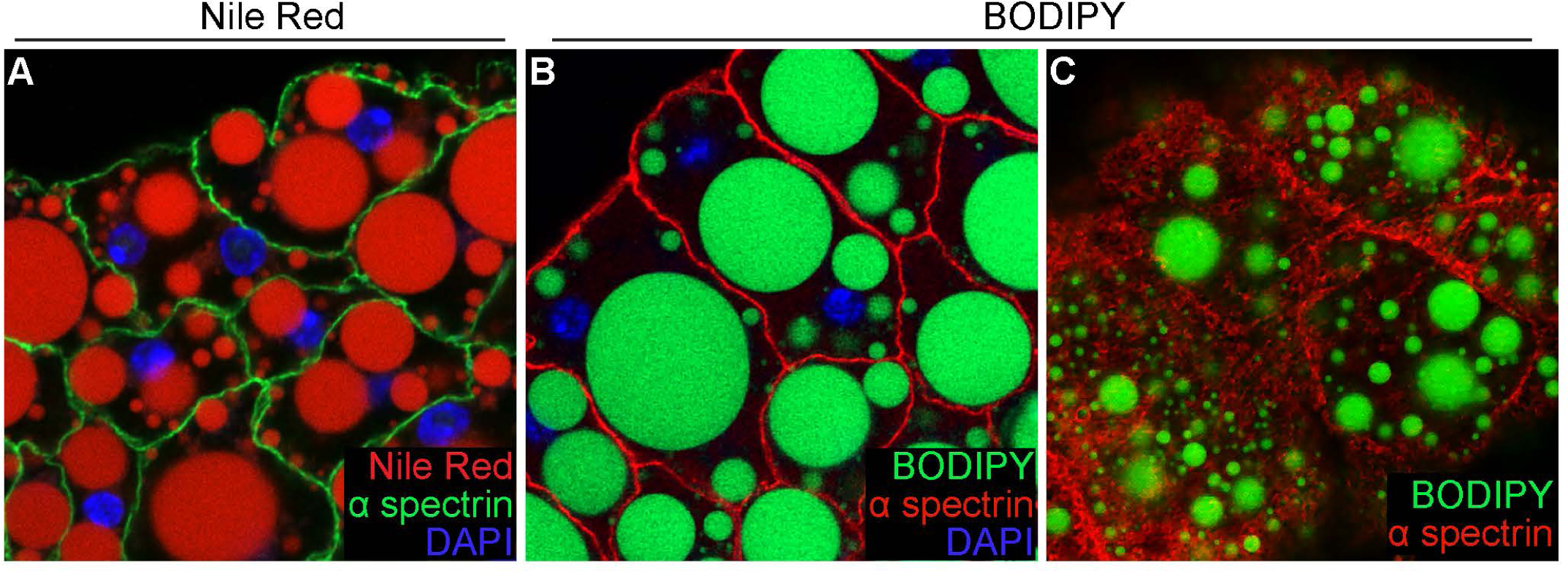
Fluorescent labeling of lipid droplets in adult adipocytes. (A) Adult adipocytes with lipid droplets labeled with (A) Nile Red (red) or (B,C) BODIPY (green). Cell membranes labeled with anti-alpha spectrin (green in A, and red in B and C) (DAPI, blue). (A,B) Single-slice confocal images taken at a middle plane of the fat body sample. (C) Single-slice confocal image taken at a surface plane of the fat body sample.

## DISCUSSION

Immunofluorescence techniques have been routinely used to examine the *Drosophila* fat body in both larvae and adults (Bradshaw et al., 2024; Jia et al., 2014; Nelliot et al., 2006; Scopelliti et al., 2019). While optical clearing methods can overcome imaging challenges posed by the adult insect cuticle (Pende et al., 2018), the delipidation steps employed in clearing protocols (Yu et al., 2021) defeats the purpose of examining lipid rich tissues like the fat body. Therefore, much of the cellular analysis of adipose tissue in *Drosophila* has focused on the larval fat body. An important step in optimizing whole-mount immunofluorescence of adult adipose tissue was to determine if abdominal carcasses made stationary by being pinned down could withstand the multi-step immunostaining process with minimal tissue damage. We find that this method of handling abdominal carcasses reduced adipose tissue loss compared to our original handling of free-floating isolated abdominal carcasses. Isolated and pinned adult fat bodies showed the expected polygonal adipocyte morphology with characteristic lipid droplet enrichment, as well as proper localization of alpha-spectrin, DE-cadherin, and filamentous actin at the cell membrane and alpha-tubulin throughout the cytoplasm. With this updated protocol, we provide a more consistent and robust approach to assess cellular morphology, protein localization, and organelle dynamics in the *Drosophila* adult fat body.

While we have identified standard conditions for fixation and permeabilization type, concentration, and time, these should serve as starting points for optimizing immunostaining protocols tailored to specific transgenic reporters and antibodies being used. For example, immunolabeling quality for DE-cadherin was sensitive to detergent type and concentration, while alpha-spectrin, alpha-tubulin, and F-actin was not (Fig. 3). This suggests that the permeabilization step should receive special attention, particularly for labeling transmembrane proteins, like DE-cadherin (Shapiro and Weis, 2009), for which epitope access may be impacted by the detergent used. Thus, several steps in an immunofluorescence protocol should be considered in advance, including cell or tissue handling, fixative, permeabilization, antigen retrieval, and primary and secondary antibody concentration and incubation time (Im et al., 2019).

The protocol described here, in conjunction with fluorescence microscopy, can be used to examine adipose tissue cellular biology during homeostasis as well as in response to genetic manipulations and/or changes in organismal physiology. In mammals, age, diet, sex, exercise, and hormones influence fat body distribution, adipocyte proliferation, differentiation, and growth, adipose tissue browning and beiging, lipid storage, metabolic activity, and gene expression (Boulet et al., 2022; Farris et al., 2024; Lu et al., 2023; Naftaly et al., 2022; Pérez et al., 2016; Stanford and Goodyear, 2016; Zhao and Yue, 2024). Similarly, *Drosophila* fat body cellular morphology, mass, molecular signature, as well as energy storage and endocrine roles respond to aging, dietary input, physical activity, and sexual dimorphism (Diaz et al., 2023; Huang et al., 2022; Lu et al., 2023; Matsuoka et al., 2017; Song et al., 2023; Wat et al., 2020; Wat et al., 2021). With the incredible *Drosophila* genetic toolkit, improvements in cell biological tools like whole-mount immunofluorescence, will allow future studies to unravel the complex molecular mechanisms that mediate adipose tissue function; thus shedding light on what goes awry during pathophysiological conditions such as premature aging and diet-induced obesity.

## MATERIALS and METHODS

### *Drosophila* strains and culture conditions

*Drosophila melanogaster* stocks were maintained at 22-25°C on standard medium containing cornmeal, molasses, yeast, and agar (Archon Scientific). Fat body samples were obtained from adult female flies fed the standard medium supplemented with wet yeast paste for one day. The *Drosophila* line *cg-Gal4; UAS-myr-RFP* (stock number 63147; *w*; P{Cg-GAL4.A}2, P{UAS-myr-mRFP}1/CyO; P{UAS-GFP.RNAi.R}142*) was obtained from the Bloomington Drosophila Stock Center (https://bdsc.indiana.edu). The previously described *FB-Gal4; UAS-GFP.nls* (*FB-Gal4, UAS-nucGFP/SM6; tub-Gal80ts/TM6b*) was used (Armstrong et al., 2014; Grönke et al., 2003). *FB-Gal4, UAS.nls* females were maintained at 22-25°C then switched to 29°C, the restrictive temperature for Gal80^ts^, to induce transgene expression prior to dissection. Females of genotypes *w^1118^* and *cg-Gal4, UAS-myr-mRFP* were maintained at 22-25°C prior to dissection.

### Pinned abdominal carcass dissection

All tissues were dissected in 1X PBS using ultra-fine forceps. Following separation of the head and thorax, abdomens were submerged in 1X PBS. A tear along the ventral anterior-posterior axis was made and internal organs removed to obtain abdominal carcasses (i.e., empty abdomens). For optimal pinning of the abdominal carcass, the last posterior abdominal segment was removed. Four 0.10 mm Austerlitz insect pins (Roboz Surgical Instrument Co.) were used to anchor each corner of the abdominal carcass to the surface of tissue culture plates (Corning Life Sciences) coated with Sylgard® 184 silicone elastomer (DOW Chemical).

### Immunostaining and confocal microscopy

All solutions used throughout the immunostaining protocol were added to and removed from tissue culture wellss. For fixation type, concentration, and timing details, see the main text. Briefly, after fixation, abdominal carcasses were washed twice for 15 minutes at room temperature in the indicated percentage of Triton X-100 or Tween 20 diluted in 1X PBS, incubated in blocking solution (5% Bovine Serum Albumin, 5% Normal Goat Serum, and Triton X-100 or Tween 20) for 3 hours at room temperature or overnight at 4°C. Abdominal carcasses were incubated overnight at 4°C in the following primary antibodies diluted in the appropriate blocking solution: mouse anti-alpha spectrin (3A9, DSHB; 3.0 ug/ml); rat anti-DE-cadherin (DCAD2, DSHB; 3.0 ug/ml); and mouse anti-alpha tubulin (12G10, DSHB; 3.0 ug/ml). Following three 15-minute washes in 1X PBS containing appropriately diluted detergent, abdominal carcasses were incubated in AlexaFluor conjugated secondary antibodies (goat anti-mouse 488, goat anti-mouse 568, or goat anti-rat 488, ThermoFisher; 1:250) for two hours at room temperature and protected from light.

For visualization of F-actin, 4% paraformaldehyde fixed tissues were washed once with 0.1% Triton X-100 in PBS before a two-hour incubation with AlexaFluor 647 Phalloidin (ThermoFisher; 1:1000) diluted in blocking solution.

For visualization of lipid droplets, alpha spectrin immunostained fat bodies were washed once with 0.1% Triton X-100 in PBS before a 30-minute incubation with BODIPY 505/515 (ThermoFisher; 25 ng/ml) diluted in deionized water or Nile Red (Sigma; 25 ng/ml) diluted in 50% glycerol. All samples were washed once prior to mounting in Vectashield containing DAPI (Vector Labs).

Fat body samples mounted on slides were stored at 4°C in the dark and imaged within 24 hours of staining. Images were acquired using the 40X oil objective on a Zeiss LSM 800 confocal microscope equipped with 2.6ZEN software.

## ACKNOWLEDGEMENTS

We thank Elizabeth Thames (Twiss Lab), Shannon Davis, and Dan Speiser for tools and reagents that allowed us to conduct initial protocol modifications. The monoclonal antibodies, anti-alpha spectrin (deposited by Branton, D. and Dubreuil, R.), anti-DCAD2 (deposited by Uemura, T.), and anti-alpha tubulin (deposited by Frankel, J. and Nelsen, E.M.) were obtained from the Developmental Studies Hybridoma Bank, created by the NICHD of the NIH and maintained at The University of Iowa, Department of Biology, Iowa City, IA 52242. Stocks obtained from the Bloomington Drosophila Stock Center (NIH P40OD018537) were used in this study. We also thank members of the Armstrong Laboratory for critical reading of the manuscript.

